# Spontaneous and ART-induced large offspring syndrome: similarities and differences in DNA methylome

**DOI:** 10.1101/2022.02.07.479430

**Authors:** Yahan Li, Jordana Sena Lopes, Pilar Coy Fuster, Rocío Melissa Rivera

## Abstract

Large/abnormal offspring syndrome (LOS/AOS) is a congenital overgrowth syndrome reported in ruminants produced by assisted reproduction (ART-LOS) which exhibit global disruption of the epigenome and transcriptome. LOS/AOS shares phenotypes and epigenotypes with the human congenital overgrowth condition Beckwith-Wiedemann syndrome. We have reported that LOS occurs spontaneously (SLOS), however, to date, no study has been conducted to determine if SLOS has the same methylome epimutations as ART-LOS. In this study, we performed whole genome bisulfite sequencing to examine global DNA methylation in SLOS and ART-LOS tissues. We observed unique patterns of global distribution of differentially methylated regions (DMRs) over different genomic contexts, such as promoters, CpG islands and surrounding regions, and repetitive sequences from different treatment groups. In addition, we identified hundreds of LOS-vulnerable DMRs across different cattle breeds when analyzing four idependent LOS experiments together. Specifically, there are 25 highly vulnerable DMRs that could potentially serve as molecular markers for the diagnosis of LOS, including at the promoters of *DMRT2* and *TBX18*, at the imprinted gene bodies of *IGF2R, PRDM8*, and *BLCAP/NNAT*, and at multiple CpG islands. We also observed tissue-specific DNA methylation patterns between muscle and blood, and conservation of ART-induced DNA methylation changes between muscle and blood. We conclude that as in ART-LOS, alterations of the epigenome are involved in the etiology of SLOS. In addition, SLOS and ART-LOS share some similarities in methylome epimutations.

## Introduction

Large/abnormal offspring syndrome (LOS/AOS) is a congenital overgrowth syndrome that has been reported in ruminants ^1,2^. Frequently observed features include macrosomia, macroglossia, umbilical hernia, organomegaly, placentomegaly, hydrallantois, increased gestation length, and increased dystocia rate ^3–13^. Most of the LOS reported in the literature have involved the use of assisted reproductive technologies (ART) ^3–13^; however, we recently reported that LOS can occur spontaneously ^2,14^, a phenomenon that in some cases may have been incorrectly ascribed to the sire’s genetics ^15,16^. Currently, there is a lack of documented incidence for both spontaneous LOS (SLOS) and ART associated LOS (ART-LOS) from the industry, although those experiencing them in their farm or practice incur steep financial losses ^2^.

We and others have reported that ART-LOS is an epigenetic disorder ^17,18^ with global alterations of transcriptome and methylome, changes in chromosomal architecture, and loss-of-imprinting at multiple imprinted domains including *IGF2R, KCNQ1, IGF2, PLAGL1, PEG3*, and *DLK1* ^3,4,17–23^. Although we recently documented that LOS occurs spontaneously, at least based on phenotypes ^2,14^, no data exist to demonstrate that the spontaneous overgrowth syndrome shares epigenotype with the ART-induced LOS.

Beckwith-Wiedemann syndrome (BWS, OMIM #130650) is the most common congenital overgrowth syndrome in humans. The incidence of BWS is approximately 1 in 10,340 live births and children conceived with the use of ART have a 10.7 relative risk of suffering from BWS ^24,25^. Clinical features frequently observed in BWS include macrosomia, macroglossia, abdominal wall defects (umbilical hernia/exomphalos), lateralized overgrowth, increased tumor incidence, hyperinsulinism, facial naevus simplex, ear malformation, organomegaly, and placentomegaly ^26^. Molecular defects found in BWS include global alteration of transcriptome and methylome, changes of chromosomal architectures, loss-of-imprinting at imprinted domains including *IGF2, KCNQ1, IGF2R, PLAGL1, PEG3, PEG10, GRB10, MEST, DLK1, IGF1R*, and *GNAS* ^22,26–32^. In addition, a subset of BWS are the result of secondary epimutations (genetic defects) which result in loss-of-imprinting ^33,34^. We have shown that ART-LOS shares phenotypes and molecular aberrations with BWS ^22,26^.

Given the phenotypic similarities between the spontaneous and the ART-induced syndromes we hypothesized that SLOS has similar methylome epimutations as ART-LOS. In this study we performed whole genome bisulfite sequencing in tissues of control, SLOS and ART-LOS to identify conserved signatures of this syndrome. We identified 25 highly vulnerable DMRs that could potentially serve as molecular markers for the diagnosis of LOS, including at the promoters of *DMRT2* and *TBX18*, at the imprinted gene bodies of *IGF2R, PRDM8*, and *BLCAP/NNAT*, and at multiple CpG islands. We conclude that as in ART-LOS, alterations of the epigenome are involved in the etiology of SLOS. In addition, SLOS and ART-LOS share some similarities in methylome epimutations.

## Results

### Animal information and phenotypes

In total, 26 animals were included in this study and were assigned to different groups (Table 1). The US_Control group contains three AI conceived Holstein breed neonate calves of average weight and with no clinical abnormalities and serves as control for other animals from the United States. The US_SLOS group contains eight SLOS calves found in the United States and the observable phenotypic abnormalities include macrosomia, macroglossia, and abdominal wall defects (Table 1 and Figure 1). The dam, sire, and sibling of US_SLOS_#6 showed no clinical abnormalities and are included for analyses to determine whether there exist inheritable methylation-specific causal effects of LOS or not.

**Figure 1.**
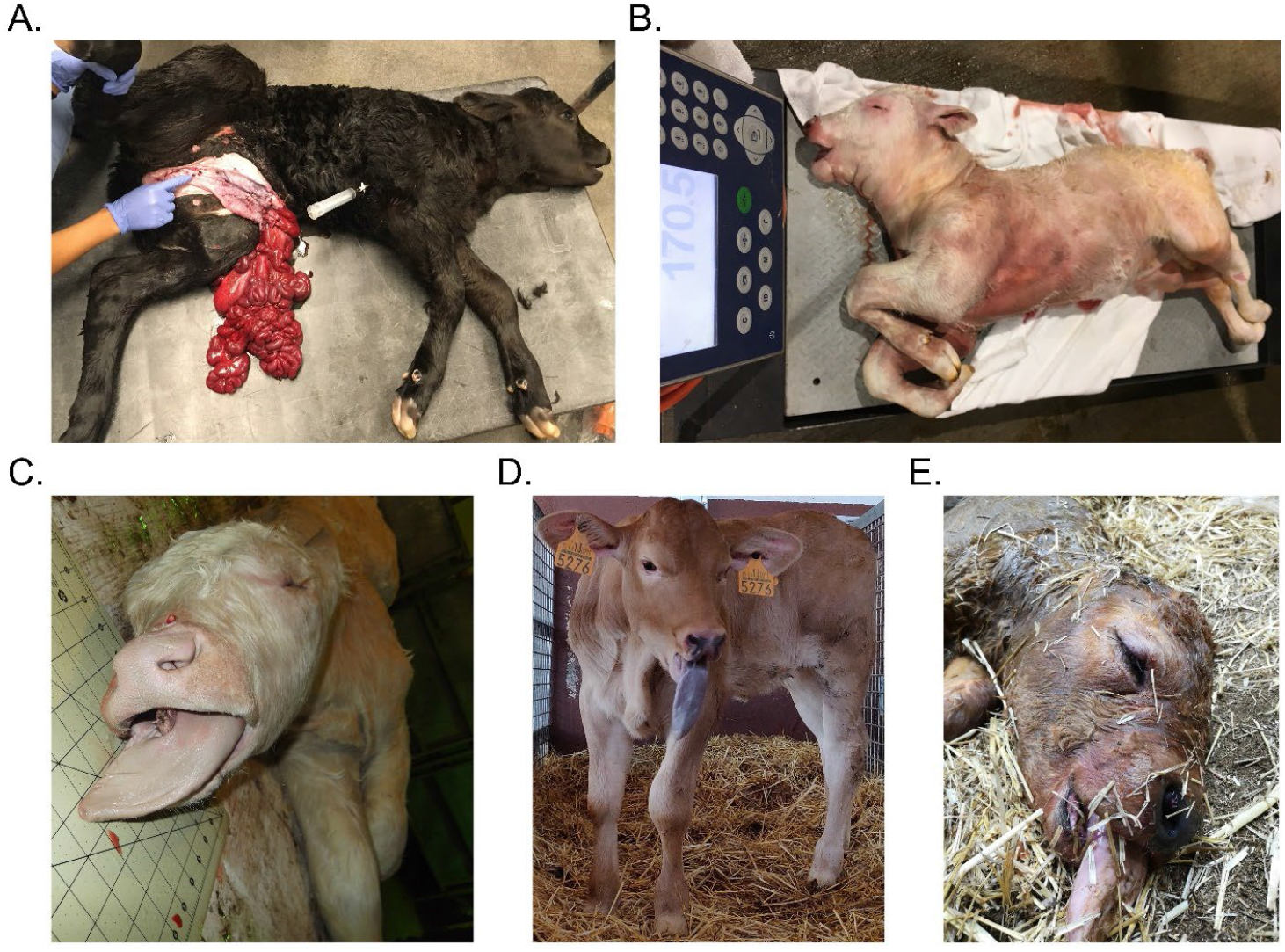
Example of phenotypic abnormalities of SLOS and ART-LOS calves. (A) Abdominal wall defect of US_SLOS_#5 (Angus breed). This spontaneous AOS calf was born alive and had to be euthanized due to the body wall malformation. (B and C) Macrosomia and macroglossia of US_SLOS_#6 (Charolais breed). This stillborn calf was ~77 Kg at birth. The average weight for calves of this breed is ~ 36 Kg. (D and E) Macroglossia of ES_ART_#2 and of stillborn ES_RF_necropsy_#1 (Asturian Valley x Limousin crossbred), respectively.

**Table 1.**
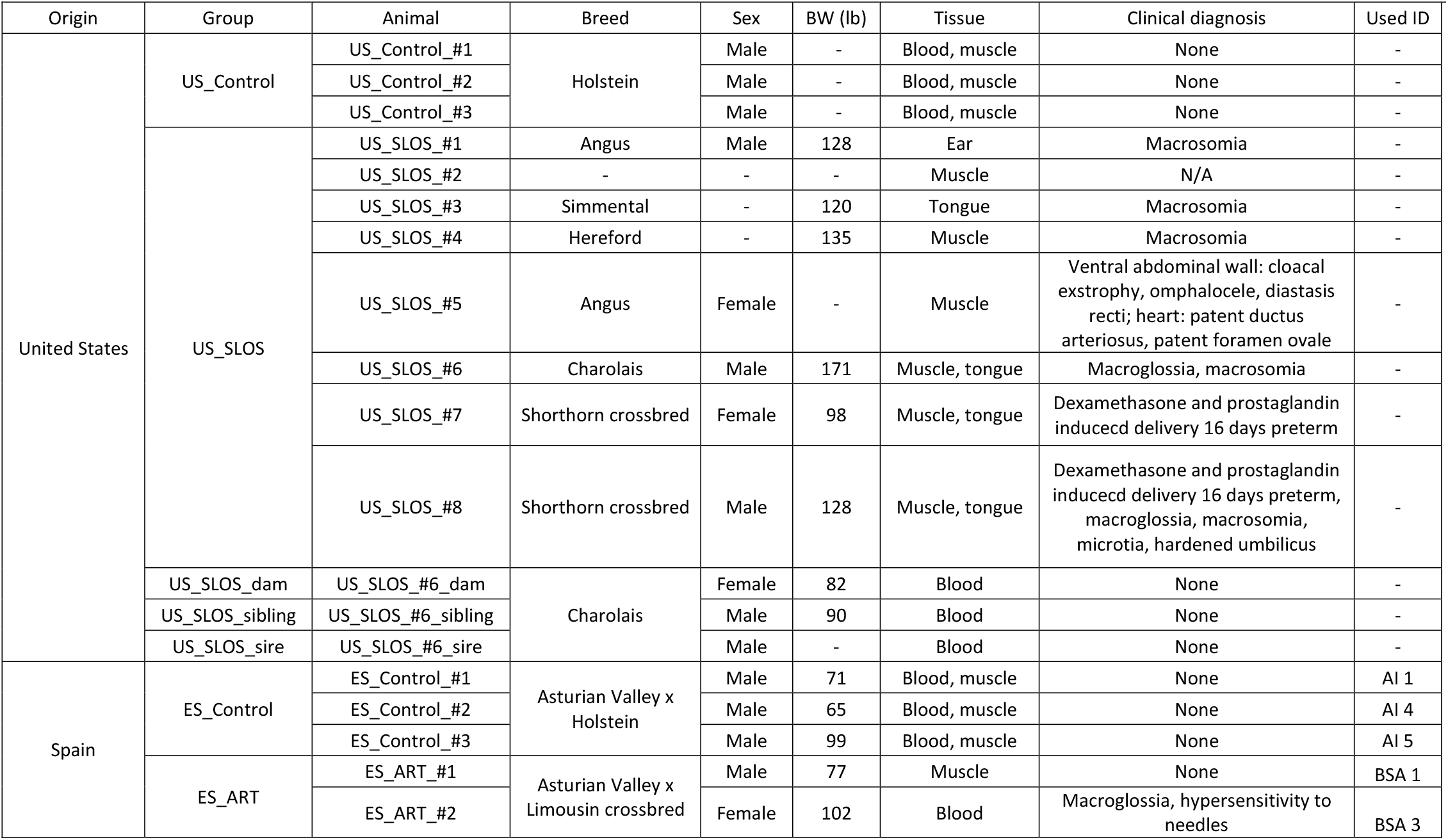
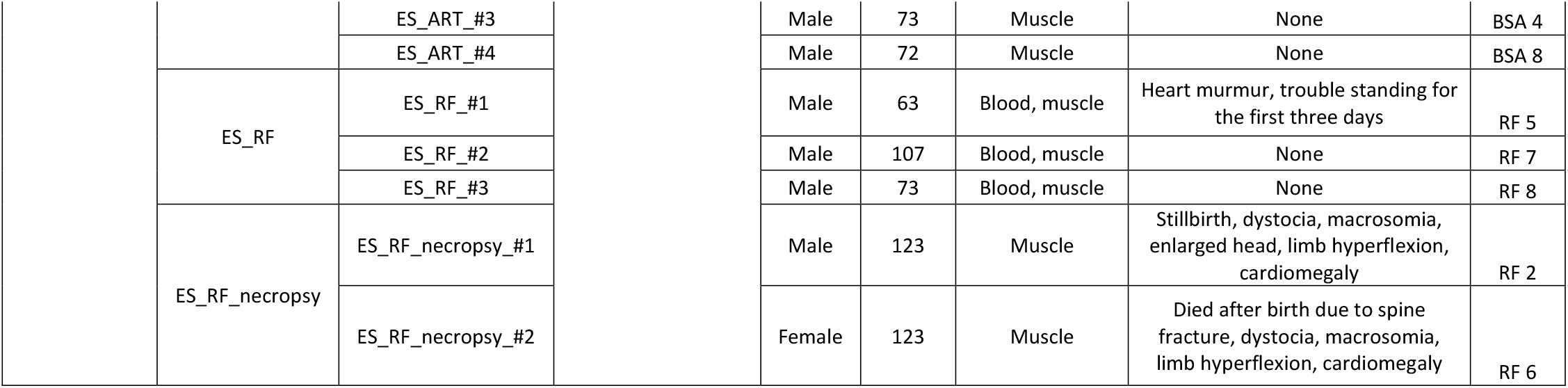
Information of calves in this study. BW = birth weight. Used ID = animal ID used in previous publications.

The ES_Control group contains three AI conceived calves with no clinical abnormalities identified and serves as control for other animals from Spain. The ES_ART group contains four ART conceived calves with no clinical abnormalities except one had macroglossia (Figure 1). The ES_RF group contains three calves conceived by ART supplemented reproductive fluids with one having some clinical abnormalities. Last, the ES_RF_necropsy group contains two dead calves from the ES_RF group with typical LOS clinical features.

### Genomic context of differentially methylated regions in SLOS calves

Whole genome bisulfite sequencing (WGBS) identified 2839 differentially methylated regions (DMRs) in US_SLOS muscle samples when compared with US_Control samples, namely US_SLOS_muscle_DMR, and ~ 66% of them were hypomethylated (Figure 2. A-C and Table S1. A). Hypomethylated DMRs more frequently occur within promoters, CpG islands, CpG shores, predicted CTCF binding sites, and repetitive sequences than hypermethylated DMRs (Figure 2. A-B). The observed frequencies of DMRs were higher than the expected frequencies at promoters, CpG islands, CpG shores, and predicted CTCF binding sites, but lower within gene bodies (both exons and introns; Figure 2. C). Of note, in this study we only included the promoters of protein coding genes and long non-coding RNAs (lncRNAs) since the location of promoters for small ncRNA are not well characterized in bovine ^35^. We also compared the DMRs identified here with those previously published for ART-LOS skeletal muscle and skin fibroblast cells ^17,23^ and identified an overlap of 22 and 134 DMRs, respectively (Figure 2. A-B). Due to the lack of proper control samples for tongue and ear tissues, a separate comparison was conducted by combining US_SLOS muscle, ear, and tongue samples and compared them to US_Control muscle samples. Similar results were observed as the muscle comparison (Figure S1. A-C and Table S1.B).

**Figure 2.**
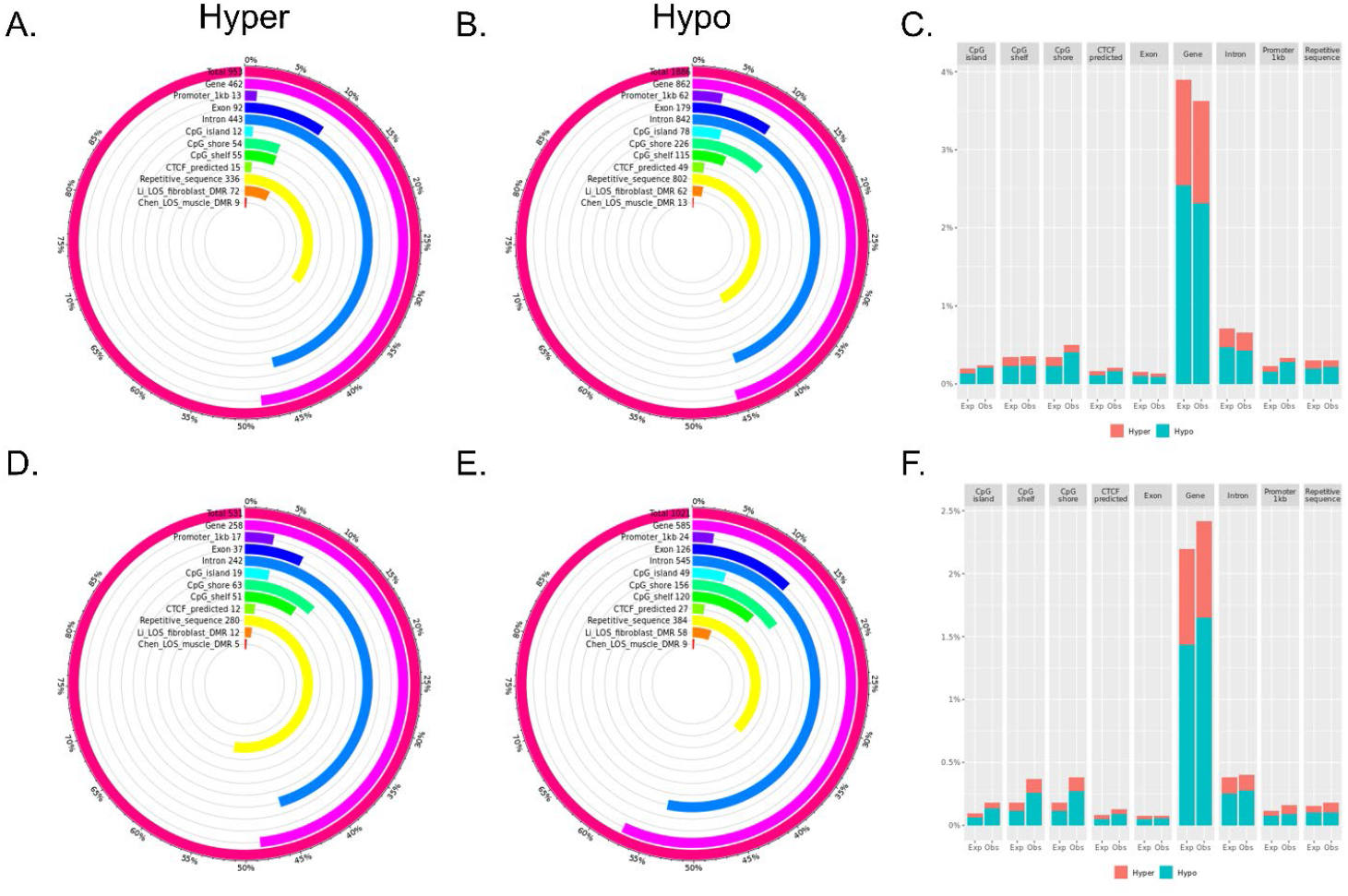
Distribution of LOS associated differentially methylated regions (DMRs) across various genomic contexts. (A-C) Muscle US_SLOS vs. US_Control DMRs. (D-F) Muscle ES_RF_necropsy vs. ES_Control DMRs. (A-B and D-E) Each figure shows the total number of DMRs in the comparison and the number and percent of the hypermethylated (hyper; A and D) and hypomethylated (hypo; B and E) DMRs over each genomic context. In addition, the figures include the number and percent of DMRs that overlap with two previous studies (Li ^23^ and Chen ^17^ for comparison purposes. (C and F) Percent of the genomic context that overlaps with DMRs. Obs = observed frequencies. Exp = expected (random frequencies obtained from shuffling DMRs across genome 1000 times).

### Genomic context of differentially methylated regions in ART-LOS calves

In total, 1,552 DMRs were identified in ES_RF_necropsy muscle samples when compared with ES_Control muscle, namely ES_RF_necropsy_muscle_DMR, and like US_SLOS_muscle_DMR, ~ 66% of DMRs were hypomethylated (Figure 2. D-F and Table S1. C). However, these hypomethylated DMRs showed a lower frequency of overlap with promoters and repetitive sequences and a higher frequency of gene body overlap when compared with hypermethylated DMRs, which is opposite with the SLOS results (Figure 2. D-E). The observed frequencies were higher than the expected frequencies in all the examined contexts except at exons (Figure 2. F).

### Vulnerable DMRs among different tissue, breeds, and developmental stages in LOS

In order to identify molecular markers for LOS, we searched for vulnerable DMRs (regardless of direction of changes: hypo or hypermethlated) observed in four independent experiments, including results of the above mentioned US_SLOS muscle and ES_RF_necropsy muscle, and our two previously published datasets, namely Li_LOS_fibroblast and Chen_LOS_muscle ^17,23^. It should be noted that the Chen_LOS_muscle raw data were reanalyzed with methods described here. In total, four DMRs were found vulnerable in all four experiments, 21 DMRs were found in three of the four experiments, and 295 DMRs were found in two of the four experiments (Figure 3 and Table S2). Overall, the DMRs found vulnerable in three to four experiments were enriched for CpG islands and CpG shores (Figure 3). The DNA methylation level and coverage for several of the LOS-vulnerable DMRs are illustrated in Figures 4 and 5 and Figures S2-S4.

**Figure 3.**
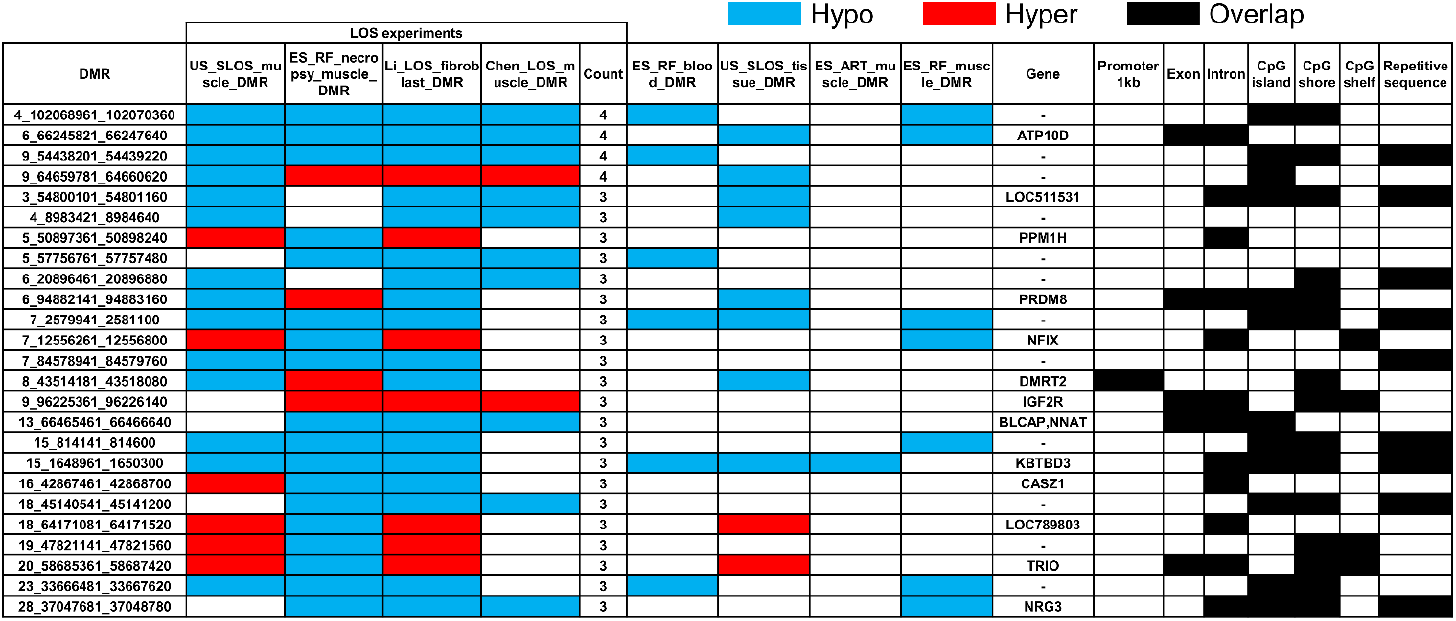
Example of LOS-associated vulnerable DMRs. Hypo = hypomethylation. Hyper = hypermethylation. DMR identifier are the positions in the bovine genome assembly ARS-UCD1.2. For complete information please refer to Table S2. DMR – differentially methylated regions.

**Figure 4.**
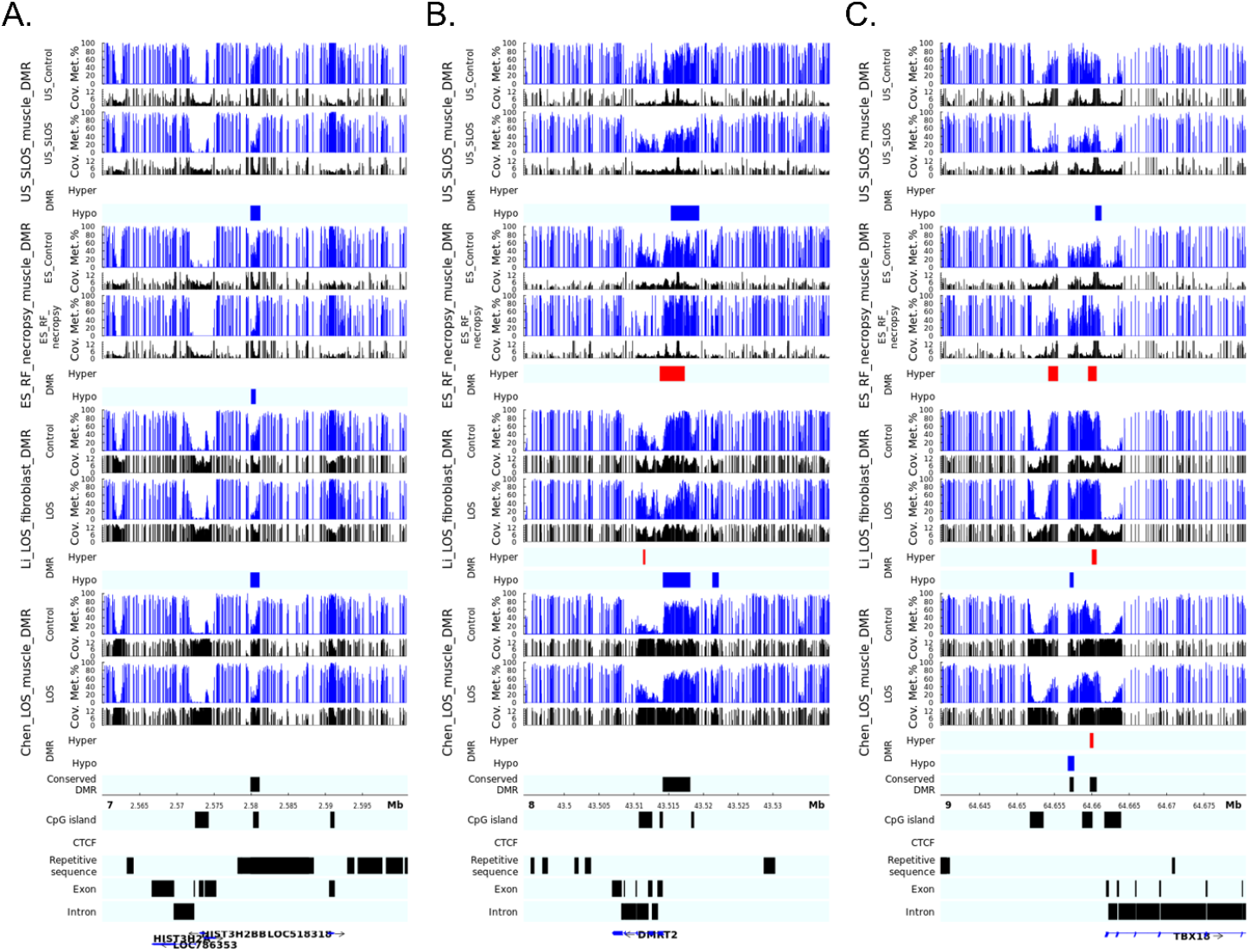
LOS-vulnerable DMRs around promoter regions. This figure shows DNA methylation level of DMRs 7_2579941_2581100 (A), 8_43514181_43518080 (B), and 9_64659781_64660620 (C) in four LOS experiments. The aforementioned numbers refer to the chromosomes and genomic position in bovine genome assembly ARS-UCD1.2. Met.% = group mean CpG methylation level in percent. Cov. = group mean CpG read coverage. Hyper = hypermethylation (red). Hypo = hypomethylation (blue).

**Figure 5.**
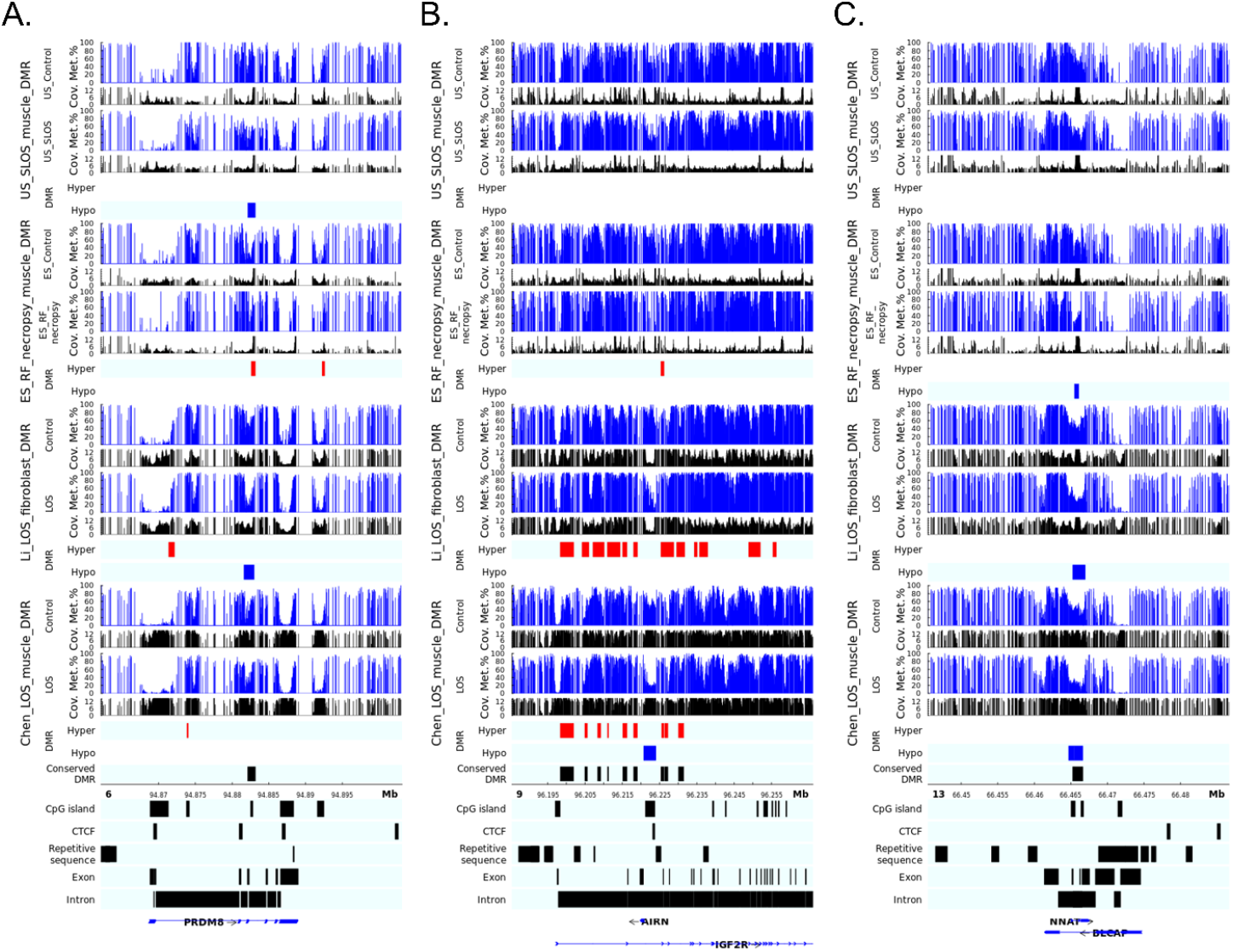
LOS-vulnerable DMRs overlapping CpG islands in body of imprinted genes. This figure shows DNA methylation level of DMRs 6_94882141_94883160 (A), 9_96225361_96226140 (B), and 13_66465461_66466640 (C) in four LOS experiments. The aforementioned numbers refer to the chromosomes and genomic position in bovine genome assembly ARS-UCD1.2. Met.% = group mean CpG methylation level in percent. Cov. = group mean CpG read coverage. Hyper = hypermethylation (red). Hypo = hypomethylation (blue).

The disruption of *IGF2R* imprinted domain has been frequently reported in LOS and BWS and we identified nine vulnerable DMRs within this domain (Figure 3 and Table S2). However, although there is ~20% and ~40% reduction in DNA methylation levels in the imprinting control region (ICR) of *IGF2R* in the US_SLOS muscle and ES_RF_necropsy muscle, respectively (Figure 5.B), hypomethylation is not reported for these samples because of the read coverage being lower than the cutoff used in this study.

### A case study for DNA methylation at LOS-associated vulnerable DMRs in a SLOS calf, its sire, dam, and full-sibling

To determine the parental impacts on LOS development, we conducted a case study for DNA methylation at LOS-associated vulnerable DMRs in US_SLOS_#6 and its sire, dam, and sibling. Due to the lack of comparable tissue samples and limited sample number, we only draw plots for visual examination of the trend of DNA methylation changes without statistical tests (Figure 6). For the 25 LOS-vulnerable DMRs shown in Figure 3, nine showed obvious differences (>10%) of DNA methylation in parental blood samples when compared to the mean of control group, including one in sire only (Figure 6. F), two in both sire and dam (Figure 6. G and H), and six in dam only (Figure 6. I to N). This higher number of conserved LOS-vulnerable DMRs in dam than in sire indicates higher proportion of maternal contribution to the SLOS development in US_SLOS_#6.

**Figure 6.**
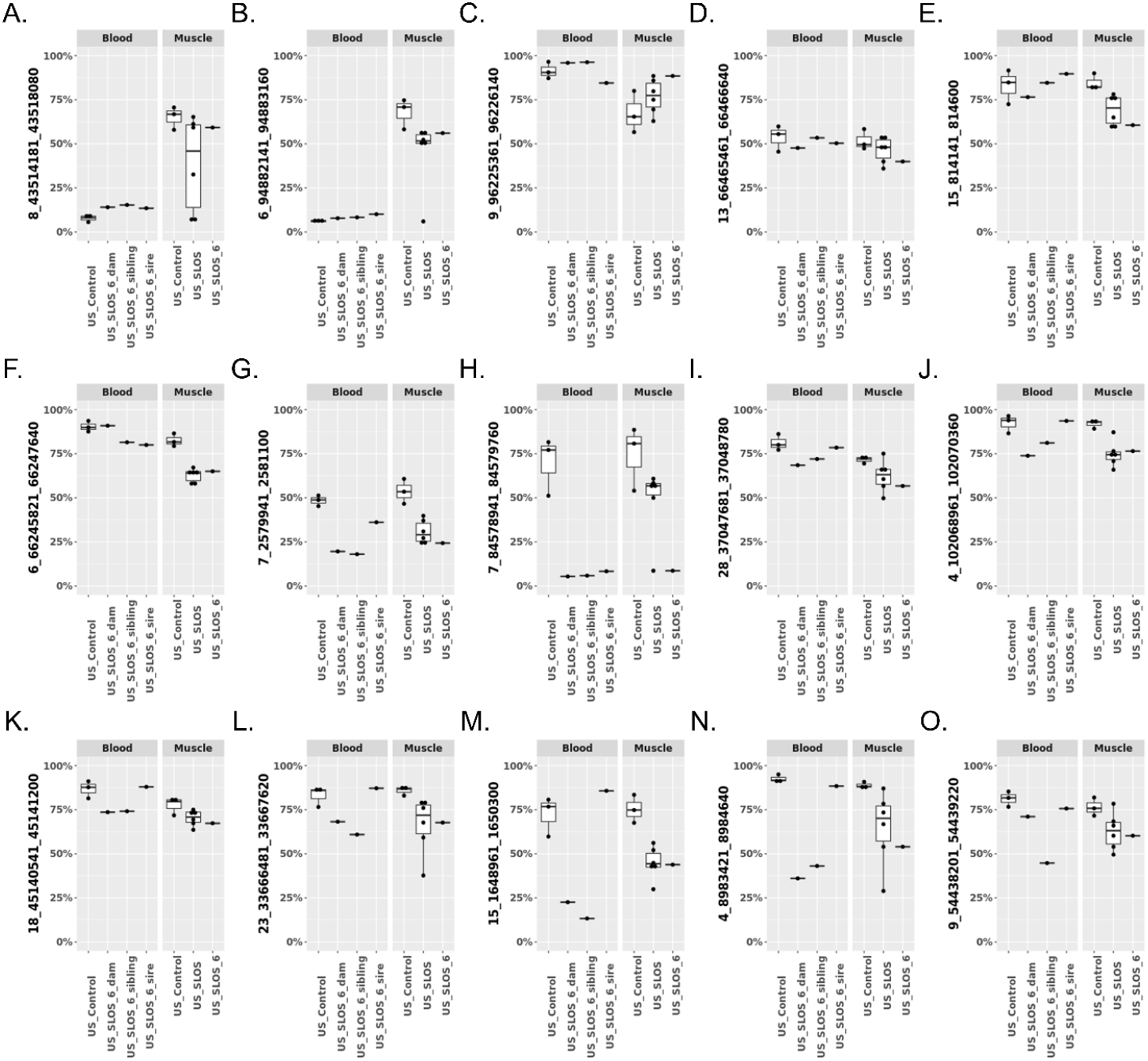
DNA methylation at LOS-vulnerable differentially methylated regions (DMRs) in US_SLOS_#6 calf and its sire, dam, and sibling. These box and dot plots show DNA methylation level (y-axis) at LOS-vulnerable DMRs without obvious differences (>10%) in parental blood samples (A-E), with obvious differences in sire only (F), both sire and dam (G-H), dam only (I-N), and sibling only (O) when compared to the mean of control group.

### Reproductive fluid supplementation partially improved methylome outcomes of ART

In total, 857 DMRs were found between ES_ART muscle and ES_Control, namely ES_ART_muscle_DMR, with a bias towards hypermethylation which is ~84% (Figure 7. A-C and Table S3. A). The observed frequencies of overlapping genomic contexts were higher than expected except at repetitive sequences (Figure 7. C). The supplementation of reproductive fluids reduced the number of DMRs (ES_RF_muscle_DMR) to 419 with about equal percent of hyper- and hypomethylation (Figure 7. D-F and Table S3. B). However, the observed frequencies of overlapping genomic contexts were still higher than expected for most examined regions (Figure 7. F). When comparing the DMRs between ES_ART_muscle_DMR and ES_RF_muscle_DMR, 62 were shared and had similar hypo or hypermethylation (Table S3. C).

**Figure 7.**
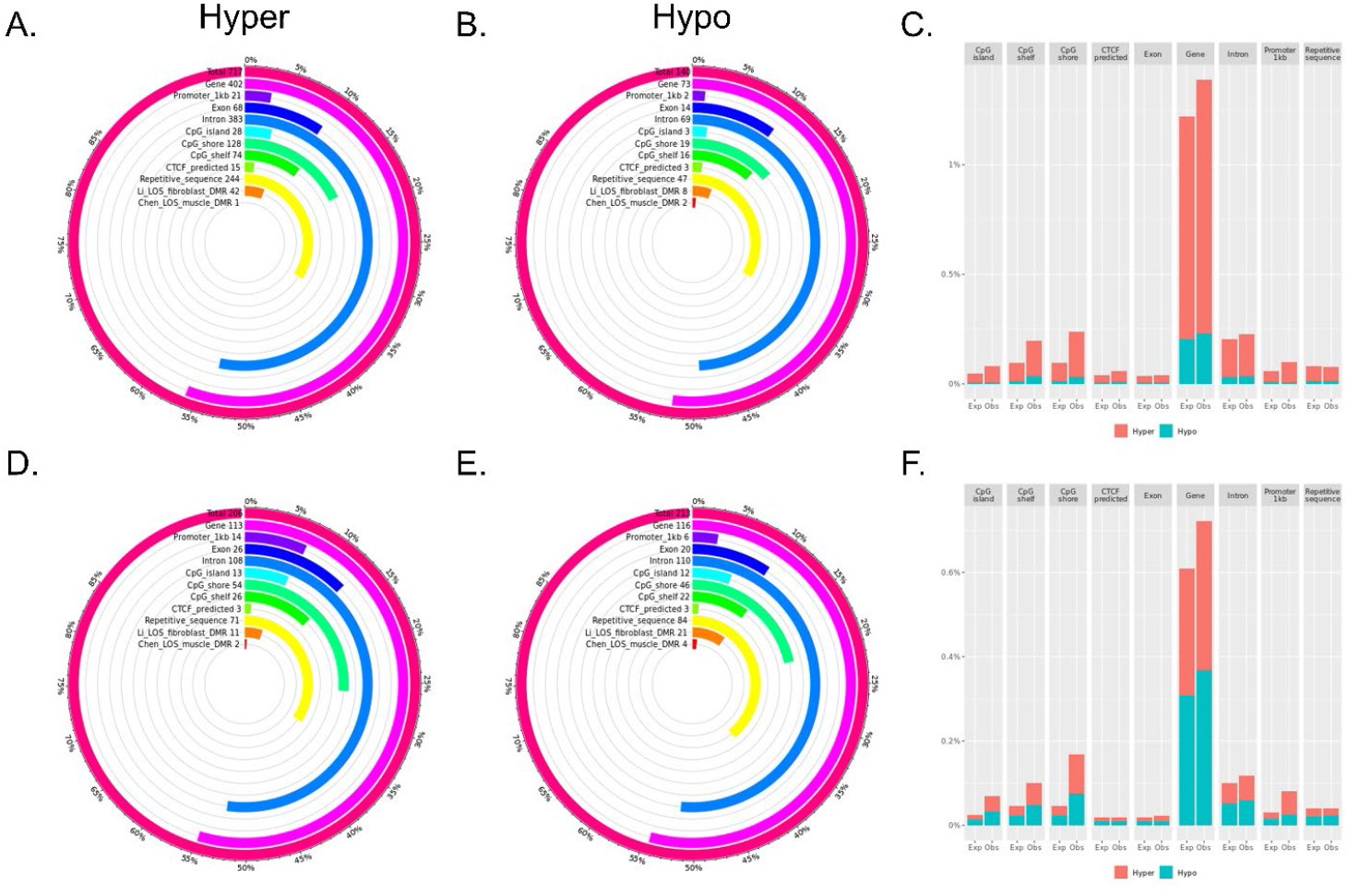
Distribution of ART associated differentially methylated regions (DMRs) across various genomic contents. (A-C) Muscle ES_ART vs. ES_Control DMRs. (D-F) Muscle ES_RF vs. ES_Control DMRs. (A-B and D-E) Each figure shows the total number of DMRs in the comparison and the number and percent of the hypermethylated (hyper; A and D) and hypomethylated (hypo; B and E) DMRs over each genomic context. In addition, the figures include the number and percent of DMRs that overlap with two previous studies (Li ^23^ and Chen ^17^ for comparison purposes. (C and F) Percent of the genomic context that overlaps with DMRs. Obs = observed frequencies. Exp = expected (random frequencies obtained from shuffling DMRs across genome 1000 times).

### Tissue specific DNA methylation pattern between muscle and leukocytes

Overall, the genome of muscle is hypomethylated when compared to leukocytes. There were a total of 25,466 and 9,961 DMRs between blood and muscle for US_Control and ES_Control, respectively, and ~90% of them were hypomethylated in muscle (Figure S5. A-F and Table S4. A-B). In addition, 5,169 DMRs were shared by these two different breeds of cattle and had similar direction of change in muscle (Table S4. C).

### Conservation of ART induced DNA methylation between muscle and leukocytes

In total, 591 DMRs were identified when comparing ES_RF blood samples to ES_Control blood, namely ES_RF_blood_DMR, and ~88% of these DMRs show hypermethylation (Figure S6. A-C and Table S5. A). This pattern was not similar to ES_RF_muscle_DMR but resembled ES_ART_muscle_DMR (Figure 7). When comparing the DMRs identified in ES_RF blood and muscle, 38 were found to be shared with 37 having the same direction in methylation change (Table S5. B). In addition, when comparing between ES_RF_blood_DMR and ES_ART_muscle_DMR, 16 DMRs were found to be shared and all had the same direction of DNA methylation (Table S5. C). Several LOS-vulnerable DMRs that show conserved DNA methylation changes between tissues in ES_RF group are illustrated in Figure 8. Additionally, similar DNA methylation changes were observed for these DMRs in the blood sample of ES_ART_#2 which has macroglossia (Figure 8).

**Figure 8.**
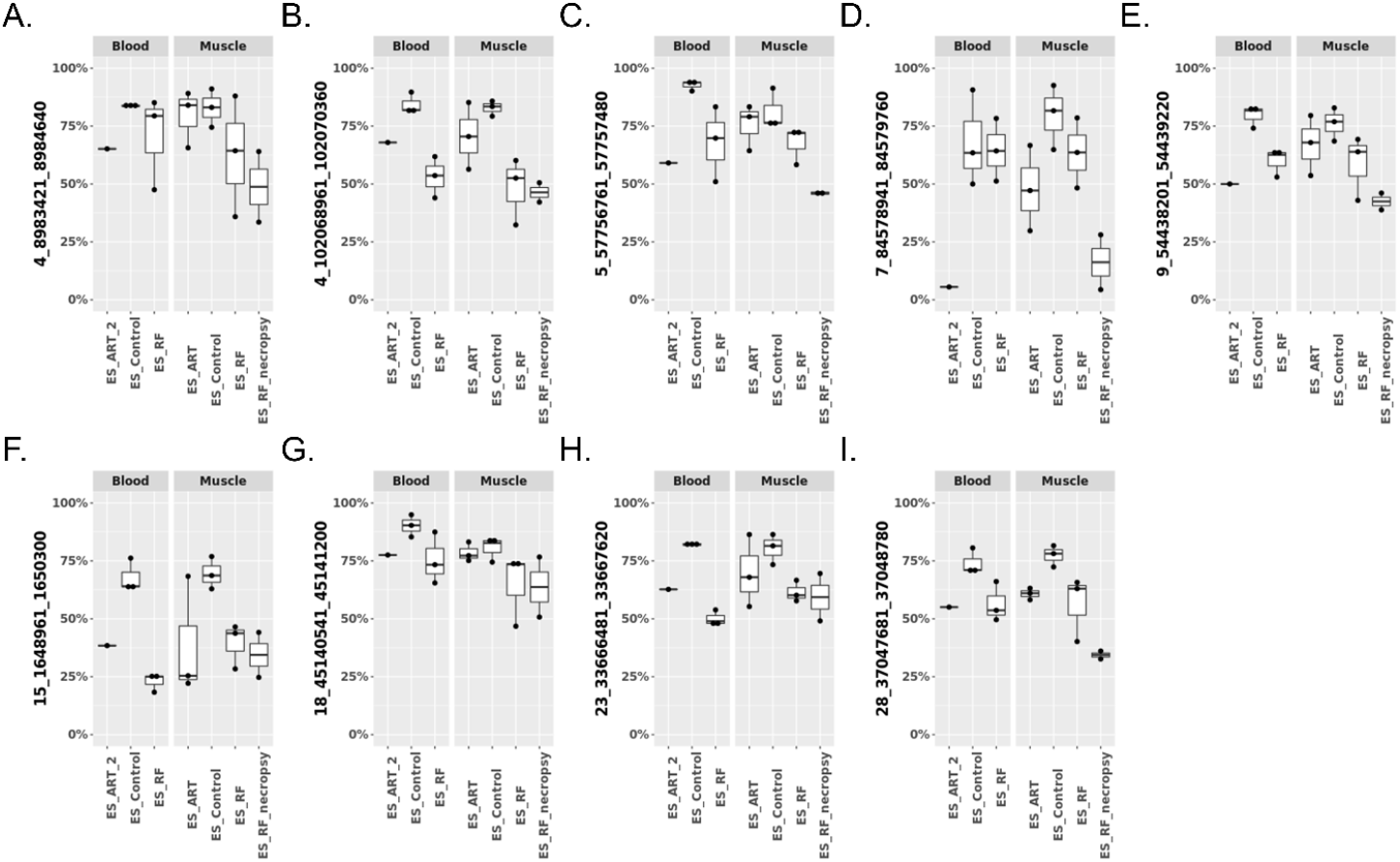
Conservation of DNA methylation at LOS-vulnerable differentially methylated regions (DMRs) between muscle and blood in ES_RF group. Y-axis = DNA methylation level. Note: ES_ART_2 is the same animal as Figure 1(D).

## Discussion

In this study, we observed typical LOS/AOS/BWS clinical abnormalities from SLOS calves, including macrosomia, macroglossia, and abdominal wall defects, and some atypical features. Spontaneous BWS (SBWS) shows no differences on the frequency of symptoms including macroglossia, hemihyperplasia, abdominal wall defects, hypoglycemia, but a significantly higher frequency of ear malformation than ART-induced BWS (ART-BWS) ^31^. In addition, SBWS patients have significantly longer gestational age and heavier birth weight than ART-BWS ^31^. Studies with larger sample size are needed to draw conclusions for these types of frequencies for SLOS.

From the analyses of DMR distribution over various genomic contexts, we saw different preference between LOS associated hyper- and hypomethylated DMRs, and similarities and differences between SLOS and ART-LOS. As a regulatory element, CpG islands and shores are enriched for enhancers in human, and the activity of enhancers is regulated by DNA methylation ^36,37^. For both US_SLOS_muscle_DMR and ES_RF_necropsy_muscle_DMR, the observed frequencies at CpG islands and shores are higher than expected, and with increased hypomethylation preference. This enrichment of hypomethylated DMRs over CpG shore resembles the observation of cancer specific DMRs in human which are associated with cell proliferation and growth ^38^. It is well known that gene expression is negatively correlated with promoter DNA methylation level ^39^. Both US_SLOS_muscle_DMR and ES_RF_necropsy_muscle_DMR showed higher frequencies overlapping promoter regions than expected, but the former had increased hypomethylation. DNA methylation level of gene bodies reflects expression level ^40^. Interestingly, US_SLOS_muscle_DMR and ES_RF_necropsy_muscle_DMR showed opposite trend of hypomethylated DMR enrichment in gene body, which suggests differences in global gene expression level.

When comparing ES_ART_muscle_DMR and ES_RF_muscle_DMR, we found the supplementation of the culture medium with reproductive fluids largely reduced the number of hypermethylated DMRs caused by ART. However, only ~15% of the DMRs of ES_RF_muscle_DMR were shared with ES_ART_muscle_DMR, indicating the supplementation of reproductive fluids also induced new changes in the methylome, which we have previously reported ^41^. Next, when adding ES_RF_blood_DMR into this comparison, we found that although the distribution pattern of ES_RF_blood_DMR resembled ES_ART_muscle_DMR, ES_RF_blood_DMR still shared more DMRs with ES_RF_muscle_DMR instead of ES_ART_muscle_DMR. This indicates that the progenitors of muscle and blood have cell type-specific response to the supplementation of reproductive fluids. Nevertheless, the shared DMRs were very consistent in the direction of changes, which suggests a that a proportion of identified DMRs could be used as diagnostic biomarkers in blood for muscle.

Several of the LOS-vulnerable DMRs were found close to the promoter of genes, including 7_2579941_2581100, 8_43514181_43518080, and 9_64659781_64660620. The hypomethylated DMR 7_2579941_2581100 resides in a region enriched for histone proteins including H2A.W histone (*H2AW/HIST3H2A*), H2B.U histone 1 (*H2BU1/HIST3H2BB*), and H3.4 histone (*H3-4/LOC518318*). However, the transcript level of these histone genes were barely detected from our previous RNA-seq results of fibroblast cells and muscle of ART-LOS fetuses ^18,23^, although that may be indicative of different developmental stages (d105 fetuses vs. newborn calves). 8_43514181_43518080 covers the promoter of protein coding gene doublesex and mab-3 related transcription factor 2 (*DMRT2*) and was detected both hypo- and hypermethylation in different experiments. *DMRT2* is a polycomb associated transcription factor and is known to be regulated by promoter DNA methylation ^42,43^. Interestingly, significant downregulation of *DMRT2* transcript was reported in muscle of ART-LOS fetuses although no DNA methylation differences detected in the corresponding samples ^18^. 9_64659781_64660620 is one of the four most frequently LOS-vulnerable DMRs mainly showing hypermethylation and located close (1.3 kb) to the transcription start site of gene T-box transcription factor 18 (T*BX18*). As a critical transcription factor during embryo development in various tissues, *TBX18* can also be regulated by promoter DNA methylation ^44,45^. The downregulation of *TBX18* transcript has been reported in muscle of ART-LOS fetuses ^18^.

We have shown that ART-LOS, like BWS is a global loss-of-imprinting disorder. Several LOS-vulnerable DMRs were found within bodies of imprinted genes and overlapped with CpG islands. 6_94882141_94883160 overlapped a CpG island within the third intron of PR/SET domain 8 (*PRDM8*). *PRDM8* is a histone methyltransferase and can inhibit cell proliferation through PI3K/AKT/mTOR signaling pathway thus functioning as a tumor suppressor ^46^. Although not completely confirmed, *PRDM8* is considered as a candidate of imprinted genes and the predicted ICR in human located in its last exon ^47,48^. The gene structure of *PRDM8* is highly conserved between human and bovine, thus this DMR overlapped CpG island is not likely to be the ICR ^48^. Accordingly, *PRDM8* was not found to be misregulated in fibroblast cells nor in muscle of ART-LOS fetuses ^18,23^. The hypomethylated DMR 13_66465461_66466640 localizes within the second intron of imprinted gene BLCAP apoptosis inducing factor (*BLCAP*) and covers most of another imprinted gene neuronatin (*NNAT*) ^49^. *BLCAP* is known as a tumor suppressor through inducing cell cycle arrest and apoptosis ^50^. *NNAT* is a proteolipid that regulates calcium channels ^51^. Increased expression of *NNAT* is often found in tumor development, including the Wilms tumor of kidney, and associated with poor outcomes of patients ^51,52^. Similarly, *NNAT* showed significant upregulation in both fibroblast cells and muscle of ART-LOS fetuses ^18,23^. Interestingly, the proposed model of imprinting regulation in human at this loci relies on CTCF binding within the second intron of *BLCAP* ^52^. However, there is no putative CTCF binding sites predicted in bovine based on vertebrate CTCF motifs, which suggests either there is undiscovered unique motif in bovine, or the mechanism of regulation is not conserved in bovine.

As previously mentioned, *IGF2R* imprinted domain contains the highest number of LOS-vulnerable DMRs, which is nine. These hypermethylated DMRs located within the first four introns of *IGF2R* surround the ICR. This ICR is the promoter of lncRNA *AIRN* and normally its methylated state on the maternal allele prevents *AIRN’s* expression and allows *IGF2R* expression ^53^. On the contrary, an unmethylated ICR on the paternal allele allows the expression of *AIRN* which silences *IGF2R* by attracting Polycomb repressive complexes to the locus ^54^. Hypomethylation of *IGF2R* ICR occurs frequently in LOS, but the low read coverage prevented us to include it in the list of vulnerable DMR although we observed similar results regardless of coverage ^17,55,56^. Compared to Chen_LOS_muscle, the decreased read coverage at *IGF2R* ICR in the other three LOS experiments is likely caused by differences in the process of sequencing library preparation. Further studies are needed to determine the reasons of this inconsistency of sequencing results of this region. DNA methylation level of gene bodies is associated with transcription frequency in a parabolic pattern that the most highly and lowly expressed genes have low level of methylation but genes with intermediate level of expression have high methylation level ^40^. This pattern matches our observation for the hypermethylated DMRs in *IGF2R* gene body and *IGF2R* transcripts were downregulated in both fibroblast cells and muscle of ART-LOS fetuses ^18,23^. For example, the *IGF2R* expression ranked 127 (top 0.7%) in the control group (~880 counts per million reads (cpm)) of fibroblast cells and decreased by ~3.5 folds to ~260 cpm in LOS group which ranked 581 (top 3%) ^23^.

Among the four most frequently LOS-vulnerable DMRs, 6_66245821_66247640 is the only one located within a gene body that does not overlap with a CpG island. This DMR covers the 20^th^ exon and surrounding 19^th^ and 20^th^ introns of ATPase phospholipid transporting 10D (*ATP10D*) gene. *ATP10D* is hypomethylated in all four LOS experiments. ATP10D functions in the modulation of high density lipoprotein and is associated with susceptibility of obesity under high fat diet in mice studies ^57^. Our previous RNA-seq results did not show misregulation of *ATP10D* transcript in either fibroblast cells or muscle of LOS fetuses ^18,23^. For the other two LOS-vulnerable DMRs found in four LOS experiments, namely 4_102068961_102070360 and 9_54438201_54439220, they always show hypomethylation at intergenic regions covering CpG islands. Further studies are needed to determine whether these DMRs serve as remote regulatory elements for gene expression.

In human, although the hierarchical cluster analyses cannot completely separate ART-BWS and SBWS groups, the two group still show different preferences of DNA methylation changes for imprinted domains including *PEG10, MEST, GNAS, PLAGL1*, and *IGF2R* ^30,31^. Similarly, we also observed different performances between SLOS and ART-LOS at some imprinted domains, including *NNAT*, IGF2R. Additionally, SBWS is associated more with genetic defects including changes of chromosomal contents and gene mutations when compared to ART-BWS ^31^. Further studies on DNA sequencing of LOS are needed to investigate if there is a genetic contribution to the susceptibility of LOS/AOS development.

Finally, we did a comparison of the methylome of a SLOS, namely US_SLOS_#6, with its relatives (dam, sire, and full-sibling) to determine if the epimutations were inherited or occurred de novo in the offspring. We identified that some of the DMRs may have been inherited through the maternal or paternal genomes, although the dam seems to contribute more to the abnormal offspring’s methylome. While some of the differences detected may be breed specific, it appears that the abnormalities in the SLOS may be partly due to the higher number of epimutations inherited from the parents as its full-sibling was born healthy and of normal size, even though it shares some inherited epimutations.

In summary, unique patterns of distribution over different genomic contexts were observed for DMRs as a result of ART, reproductive fluid supplementation of culture media, ART-LOS, and SLOS. Hundreds of LOS-vulnerable DMRs determined in this study could serve as molecular markers for the diagnosis of LOS. Further studies are needed to determine the level of conservation of these DMRs in other tissue types of LOS fetuses that could be used for early diagnosis, such as amniotic fluid. In conclusion, alterations of epigenome are involved in the etiology of SLOS with certain levels of similarities to ART induced LOS.

## Materials and Methods

All the chromosomal coordinates in this manuscript refer to bovine genome assembly ARS-UCD1.2 ^58^.

### Animal tissues

Blood and tissue samples of animals from the United States (US) and Spain (ES) were used in this study (Table 1). Control animals from the US (US_Control) were conceived by artificial insemination (AI) at the University of Missouri Foremost Dairy Research Center and sacrificed immediately upon birth by a trained Veterinarian for blood and tissue collection. The three Holstein breed neonates were male, of average birth weight and without any abnormal phenotypes. Blood samples were collected from the jugular vein using K3EDTA vacutainers (BD) and processed as described by Ortega et al. ^59^. Tissues were dissected, diced, sealed in aluminum foil pockets, snap frozen in liquid nitrogen, and stored at −80 °C.

Blood and tissue samples of stillborn SLOS animals from various parts of the US (US_SLOS) were collected by their owners, veterinarians, or our collaborators and shipped to University of Missouri. Blood samples of the dam, sire, and sibling of US_SLOS_#6 were collected by us.

Animals from Spain were generated as described previously ^60^. Briefly, the control animals (ES_Control) were conceived by AI using frozen-thawed semen from one bull (Asturian Valley breed) among synchronized cows (Holstein breed) on the day of presumptive estrus. *In vitro* produced animals were generated using slaughterhouse oocytes (crossbred Limousine and Charolais) and semen from the same bull as controls. Following fertilization, embryos were separated in two different groups: one culture group (ES_ART) composed of synthetic oviductal fluid (SOF) media supplemented with bovine serum albumin during the 7-8 days of culture and another group (ES_RF) composed of SOF media supplemented with bovine oviductal fluid (NaturARTs-BOF-EL, Embryocloud, Spain) for the first 4 days and bovine uterine fluid (NaturARTs-BUF-ML, Embryocloud) for the following days. Embryos (blastocysts and expanded blastocysts) were vitrified on day 7 or 8 of culture and stored until use. Recipients (Holstein cows) were synchronized and on day 6 to 8 after presumptive estrus, each cow received one thawed embryo. After parturition, calves were immediately assessed for general health parameters and continued to be monitored throughout their lives. Calves that did not survive parturition or died were collected for necropsy. Simultaneous blood and muscle samples were collected in two different days and calves’ age ranged between 71 and 292 days (mean age 167 days and median 138). Necropsy muscle samples do not have corresponding blood sample and the age of the calves varied between 0 (at birth) and 13 days (mean age 5 days and median 2). Blood samples were collected from the jugular or coccygeal vein (according to the size of the animal) using EDTA tube (BD vacutainer, BD, Spain) and stored at 4°C. Samples (less than 2h after collection) were then aliquoted in 300 μL and 900 μL of Tris-HCl solution was added. The content was mixed and centrifuged at 14500 x g for one minute and the supernatant discarded. The procedure was repeated twice more and the final pellet was submerged in liquid nitrogen and stored at −80°C. Muscle biopsies were performed using a semiautomatic needle (ML18160, RI.MOS., Italy). Surgical preparation prior biopsy included minor restraint of the animal, shaving of the area, cleaning, and application of local anesthesia (lidocaine). The incision on the gluteus medius was ~1cm long, enough for the biopsy needle to pass. Samples were immediately collected and placed on ice. The incision was closed, and calves were monitored for any sign of infection. Samples were transported to the laboratory (less than 2h after collection), frozen in liquid nitrogen and stored at −80 °C. Blood and muscle samples were shipped on dry ice to the University of Missouri.

### Ethics approval

US_Control animals were purchased from the University of Missouri Foremost Dairy Research Center and euthanized by veterinarians. All the animal procedures were approved by University of Missouri Animal Care and Use Committee under protocol 9455.

Animals from Spain were handled by veterinarians following the Spanish Policy for Animal Protection RD 53/2013, which meets European Union Directive 2010/63/UE on animal protection. The Ethics Committee of Animal Experimentation of the University of Murcia and the Animal Production Service of the Agriculture Department of the Region of Murcia (Spain) (ref. no. A132141002) approved the procedures performed for these animals.

### Genomic DNA extraction

Blood and tissue samples were lysed in lysis buffer (0.05 M Tris-HCl (pH 8.0), 0.1 M EDTA, and 0.5% (w/v) SDS) with proteinase K (Fisher BioReagents, BP1700) at 55°C for four hours (blood) or overnight (tissue). Genomic DNA was extracted with Phenol:Chloroform:Isoamyl Alcohol (SIGMA, P3803) following the manufacturer’s instructions. The concentration of DNA was measured by using a NanoDrop^®^ ND-1000 Spectrophotometer (Thermo Fisher Scientific) and DNA integrity was confirmed by electrophoresis on a 0.7% agarose gel. Genomic DNA samples were stored at −20°C.

### Whole genome bisulfite sequencing (WGBS) and data analyses

WGBS was conducted by CD Genomics. Information on library preparation and sequencing obtained from the company is as follows: For WGBS library preparation, 1 ug of genomic DNA was fragmented by sonication to a mean size of approximately 200-400 bp. Fragmented DNA was end-repaired, 5’-phosphorylated, 3’-dA-tailed and then ligated to methylated adapters. The methylated adapter-ligated DNAs were purified using 0.8× Agencourt AMPure XP magnetic beads and subjected to bisulfite conversion by ZYMO EZ DNA Methylation-Gold Kit (zymo). The converted DNAs were then amplified using 25 μl KAPA HiFi HotStart Uracil+ ReadyMix (2X) and 8-bp index primers with a final concentration of 1 μM each. The constructed WGBS libraries were then analyzed by Agilent 2100 Bioanalyzer and quantified by a Qubit fluorometer with Quant-iT dsDNA HS Assay Kit (Invitrogen), and finally sequenced on Illumina Hiseq X ten sequencer. 0.1-1% lambda DNA were added during the library preparation to monitor bisulfite conversion rate.

For WGBS data analyses, duplicated reads generated during PCR and sequencing were removed from raw sequencing reads using the clumpify function of BBMap 38.90 ^61^. The remaining raw reads were trimmed for adapter sequences and low quality bases using trimmomatic 0.39 ^62^ with parameters ‘ILLUMINACLIP:adapter_seq:2:30:10:1:true LEADING:20 TRAILING:20 AVGQUAL:20 MAXINFO:0:0.5’. Trimmed reads were aligned to the bovine genome using bismark 0.23.0 ^63^ with parameters ‘-X 900 --unmapped --ambiguous --non_bs_mm’. Trimmed reads were also aligned to lambda phage genome to determine bisulfite conversion rates. Samtools 1.13 ^64^ was used to convert, sort, filter, and index bam files. MarkDuplicates function of picard 2.25.5 ^65^ was used to further remove duplicated reads after alignment. Read groups were added for each samples using AddOrReplaceReadGroups function of picard. The dataset of known variants in bovine, namely ARS1.2PlusY_BQSR_v3.vcf.gz, was acquired from the 1000 bull genome project ^66^ and served as reference to identify genomic variants in WGBS data. Indel realignment was performed using RealignerTargetCreator and IndelRealigner functions of BisSNP 1.0.1 ^67^. Base quality recalibration was carried out using BisulfiteCountCovariates and BisulfiteTableRecalibration functions of BisSNP 0.82.2 since these functions are missing in version 1.0.0 and 1.0.1. Parameters used for BisulfiteCountCovariates were ‘-cov ReadGroupCovariate -cov QualityScoreCovariate - cov CycleCovariate -baqGOP 30’. Genomic variants were identified using BisSNP 1.0.1 with default setting expect that ‘-bsRate’ was changed to bisulfite conversion rate observed from lambda phage genome alignment for each sample. BisSNP identified variants were filtered by its VCFpostprocess function with parameter ‘-windSizeForSNPfilter 0’. Additionally, genomic variants were identified using BS-SNPer 1.0 ^68^ with parameters ‘-minhetfreq 0.1 --minhomfreq 0.85 --minquali 15 --mincover 5 -maxcover 1000 --minread2 2 --errorate 0.02 --mapvalue 20’. M-bias plots were generated using bismark and the first 3 bases of R1 reads and the first 4 bases of R2 reads showed biased CpG methylation level, thus these bases were excluded from downstream analyses. CpG methylation information were extracted from the bam files using bismark_methylation_extractor function of bismark with parameters ‘-p --ignore 3 --ignore_r2 4 --comprehensive --no_header -gzip --bedGraph --buffer_size 50% -- cytosine_report’. Statistical analyses were conducted using R package hummingbird ^69^ with parameter ‘minCpGs = 10, minLength = 100, maxGap = 300’ to identify differentially methylated regions (DMRs) in various comparisons. DMRs with at least 15% difference in methylation level (both gain and loss of methylation) and at least 4 mean read coverage at CpG sites were reported. The sex chromosomes were not analyzed to circumvent confounding created by X chromosome inactivation associated DNA methylation.

### Analyses of overlapping between DMRs and genomic contents

Information of gene annotation was obtained from NCBI (GCF_002263795.1_ARS-UCD1.2_genomic.gff) ^70^. Repeated and overlapped exons were merged for each gene, and introns were calculated based on merged exons. Promoters (1kb) were calculated based on transcription start sites annotation and only included protein coding genes and long non-coding RNAs. Annotation of CpG islands and repeated sequences were obtained from UCSC Genome Browser ^71^. Locations of CpG shores (flanking 2kb from CpG islands) and shelves (flanking 2-4kb from the CpG island) were calculated based on CpG island annotation. Potential CTCF binding sites were predicted as previously reported ^23^. Bedtools and custom Perl scripts were used for these analyses to identify overlapped genomic location and make tables ^72^. R package Sushi, circular, and ggplot2 were used for making figures ^73–75^.

### Sequencing data availability statement

The raw sequencing reads of WGBS used in this study are available in the GEO database with accession numbers (GSEXXX-pending).

## Acknowledgements

The work in the US was supported by Agriculture and Food Research Initiative (grant AFRI - 2018-67015-27598). We thank Astrid Roshealy Brau for technical assistance and Dr. Callum Donnelly and Mr. Stephenson for providing pictures and samples.

The work in Spain was funded by European Union, Horizon 2020 Marie Sklodowska-Curie Action (Ref. REPBIOTECH 675526), by the Spanish Ministry of Science and Innovation (MCIN/AEI/10.13039/501100011033/ and FEDER, ref. I+D+I PID2020-113366RB-I00) and Fundación Séneca, Murcia, Spain (ref. 20040/GERM/16). We thank Dr. Rafael Latorre for aiding in biopsy collection and the Physiology of Reproduction group for support with the animals.

---

**Figure S1.**
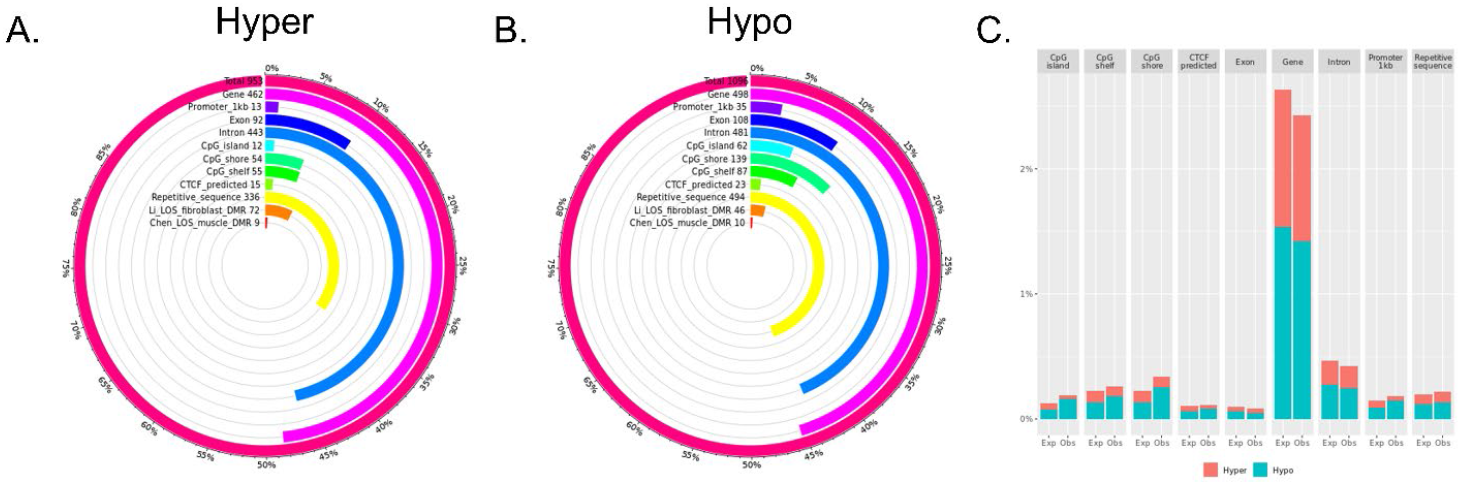
Distribution of LOS associated differentially methylated regions (DMRs) across various genomic contexts. US_SLOS tissue vs. US_Control muscle DMRs. (A-B) Each figure shows the total number of DMRs in the comparison and the number and percent of the hypermethylated (hyper; A) and hypomethylated (hypo; B) DMRs over each genomic context. In addition, the figures include the number and percent of DMRs that overlap with two previous studies (Li ^1^ and Chen ^2^ for comparison purposes. (C) Percent of the genomic context that overlaps with DMRs. Obs = observed frequencies. Exp = expected (random frequencies obtained from shuffling DMRs across genome 1000 times).

**Figure S2.**
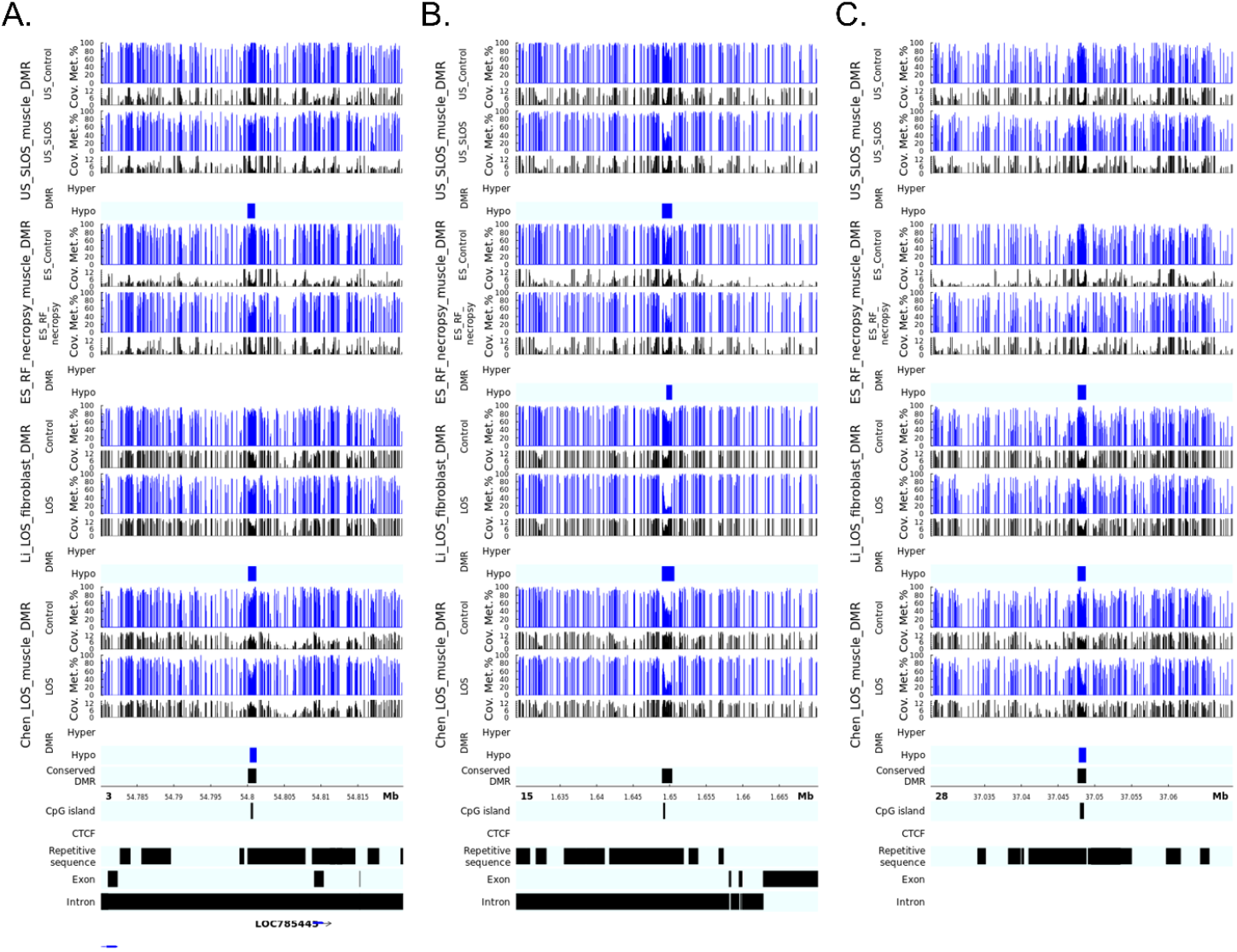
LOS-vulnerable DMRs overlapping CpG islands in gene bodies. This figure shows DNA methylation level of DMRs 3_54800101_54801160 (A), 15_1648961_1650300 (B), and 28_37047681_37048780 (C) in four LOS experiments. The aforementioned numbers refer to the chromosomes and genomic position in bovine genome assembly ARS-UCD1.2. Met.% = group mean CpG methylation level in percent. Cov. = group mean CpG read coverage. Hyper = hypermethylation (red). Hypo = hypomethylation (blue).

**Figure S3.**
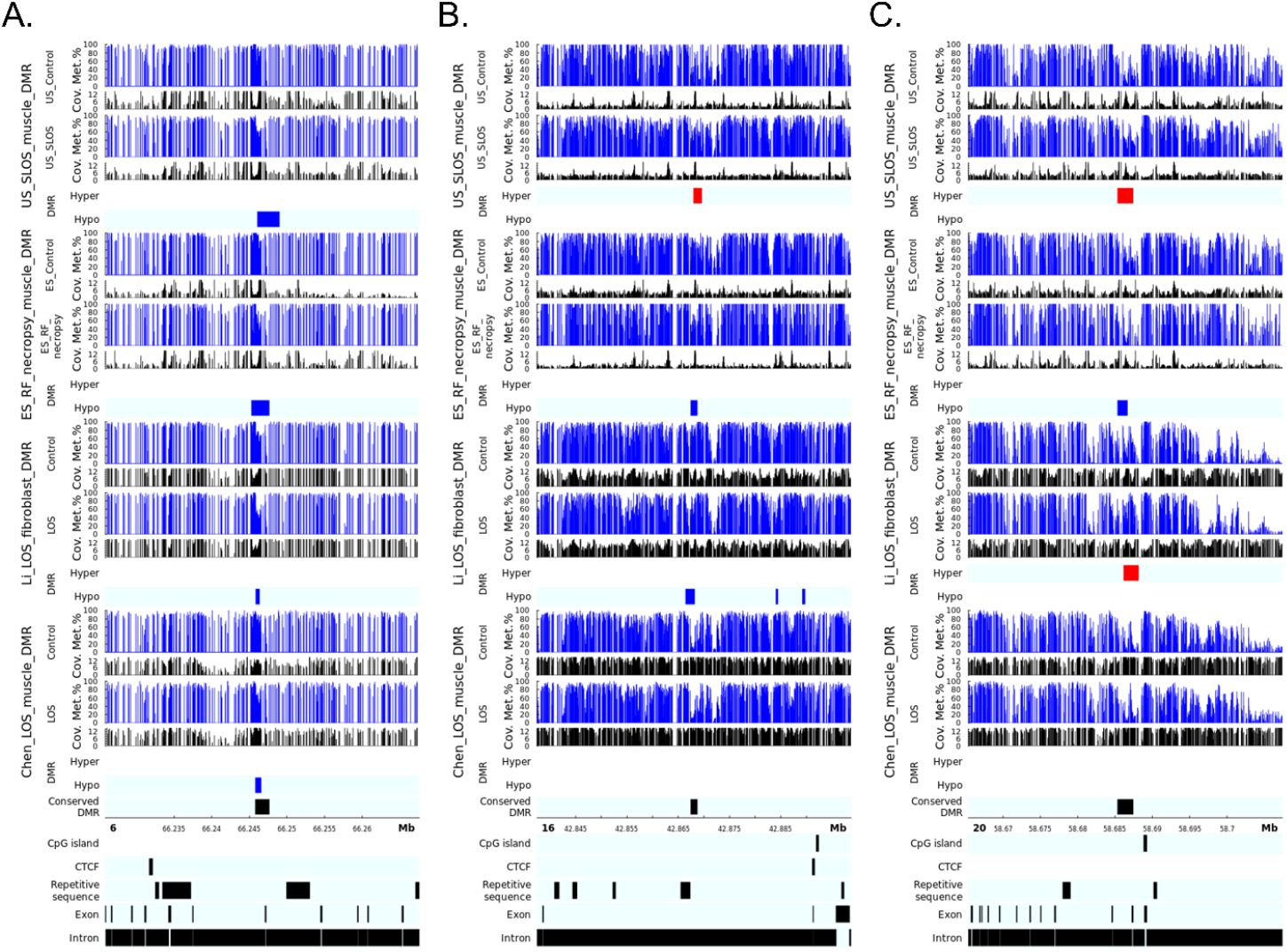
LOS-vulnerable DMRs in gene bodies. This figure shows DNA methylation level of DMRs 6_66245821_66247640 (A), 16_42867461_42868700 (B), and 20_58685361_58687420 (C) in four LOS experiments. The aforementioned numbers refer to the chromosomes and genomic position in bovine genome assembly ARS-UCD1.2. Met.% = group mean CpG methylation level in percent. Cov. = group mean CpG read coverage. Hyper = hypermethylation (red). Hypo = hypomethylation (blue).

**Figure S4.**
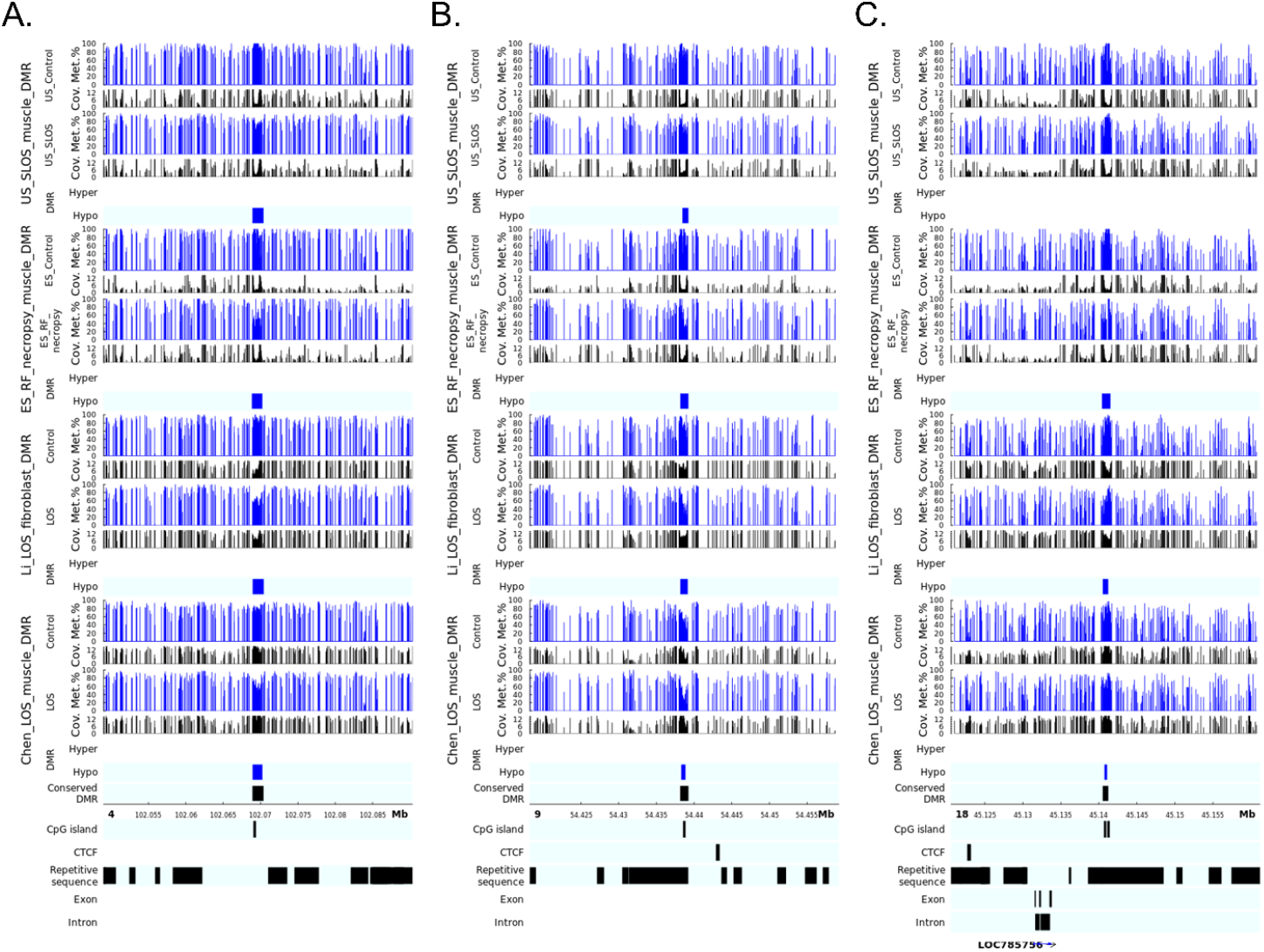
LOS-vulnerable DMRs overlapping CpG islands in intergenic regions. This figure shows DNA methylation level of DMRs 4_102068961_102070360 (A), 9_54438201_54439220 (B), and 18_45140541_45141200 (C) in four LOS experiments. The aforementioned numbers refer to the chromosomes and genomic position in bovine genome assembly ARS-UCD1.2. Met.% = group mean CpG methylation level in percent. Cov. = group mean CpG read coverage. Hyper = hypermethylation (red). Hypo = hypomethylation (blue).

**Figure S5.**
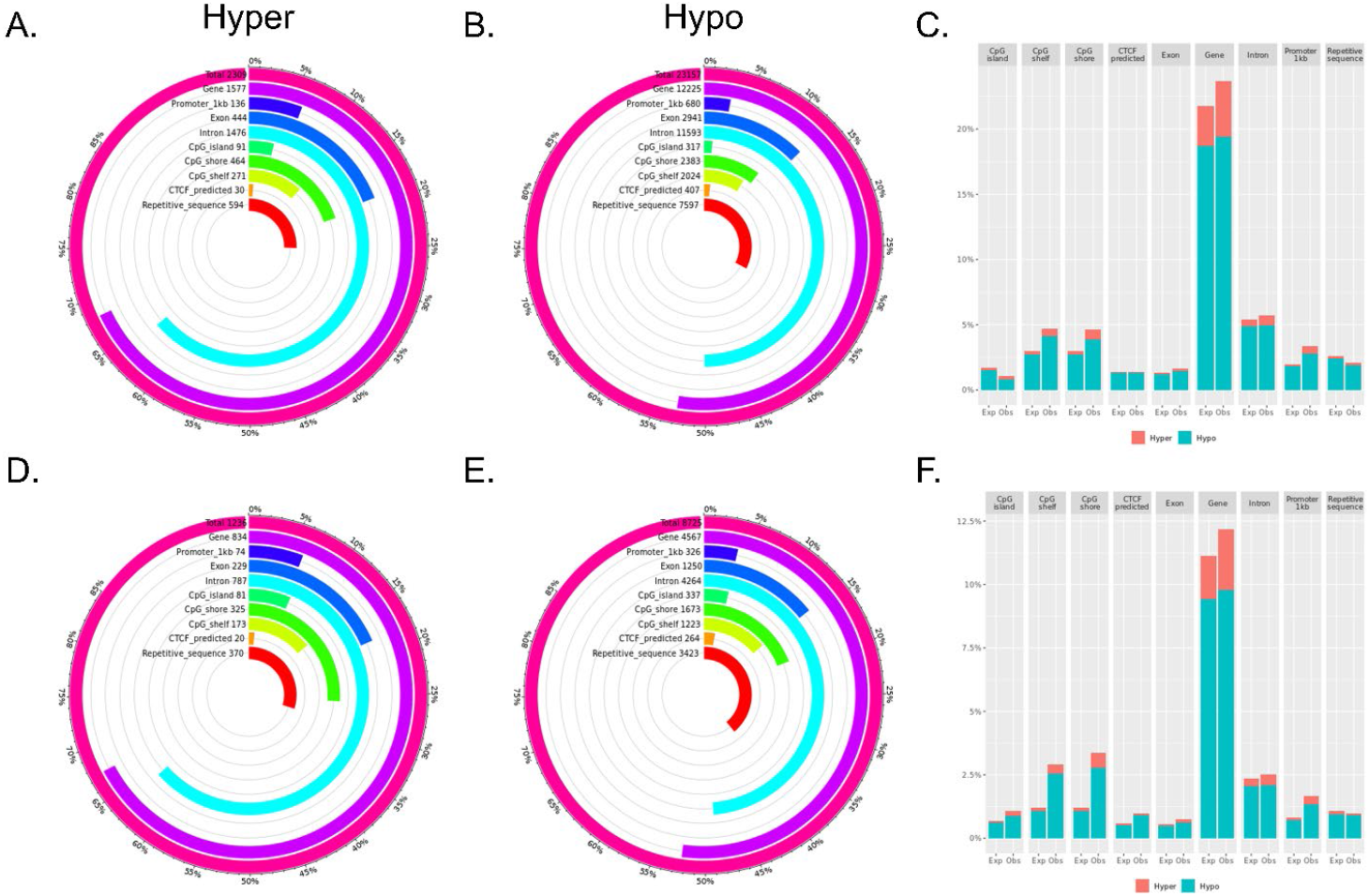
Distribution of tissue specific differentially methylated regions (DMRs) across various genomic contents. (A-C) US_Control muscle vs. blood DMRs. (D-F) ES_Control muscle vs. blood DMRs. (A-B and D-E) Each figure shows the total number of DMRs in the comparison and the number and percent of the hypermethylated (hyper; A and D) and hypomethylated (hypo; B and E) DMRs over each genomic context. (C and F) Percent of the genomic context that overlaps with DMRs. Obs = observed frequencies. Exp = expected (random frequencies obtained from shuffling DMRs across genome 1000 times).

**Figure S6.**
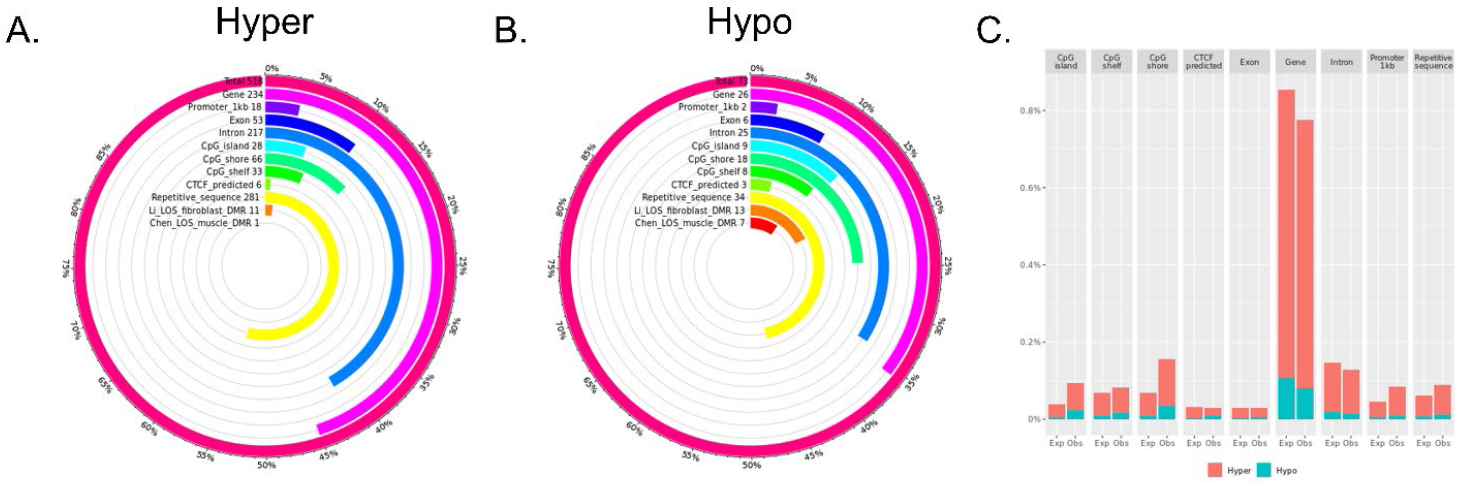
Distribution of ART associated differentially methylated regions (DMRs) across various genomic contents. Blood ES_RF vs. ES_Control DMRs. (A-B) Each figure shows the total number of DMRs in the comparison and the number and percent of the hypermethylated (hyper; A) and hypomethylated (hypo; B) DMRs over each genomic context. In addition, the figures include the number and percent of DMRs that overlap with two previous studies (Li ^1^ and Chen ^2^ for comparison purposes. (C) Percent of the genomic context that overlaps with DMRs. Obs = observed frequencies. Exp = expected (random frequencies obtained from shuffling DMRs across genome 1000 times).

